# MEGAN-LR: New algorithms allow accurate binning and easy interactive exploration of metagenomic long reads and contigs

**DOI:** 10.1101/224535

**Authors:** Daniel H. Huson, Benjamin Albrecht, Caner Bagci, Irina Bessarab, Anna Gorska, Dino Jolic, Rohan B.H. Williams

**Affiliations:** Center for Bioinformatics, University of Tübingen, Sand 14, 72076 Tübingen, Germany; Life Sciences Institute, National University of Singapore, 28 Medical Drive, Singapore 117456; Max-Planck Institute for Developmental Biology, 72076 Tübingen, Germany; Singapore Centre for Environmental Life Sciences Engineering, National University of Singapore, 28 Medical Drive, Singapore 117456; IMPRS ‘From Molecules to Organisms’, Tübingen, Germany

**Keywords:** microbiome, long reads, sequence analysis, taxonomic binning, functional binning, algorithms, software, Nanopore, PacBio

## Abstract

**Background:** There are numerous computational tools for taxonomic or functional analysis of microbiome samples, optimized to run on hundreds of millions of short, high quality sequencing reads. Programs such as MEGAN allow the user to interactively navigate these large datasets. Long read sequencing technologies continue to improve and produce increasing numbers of longer reads (of varying lengths in the range of 10k-1M bps, say), but of low quality. There is an increasing interest in using long reads in microbiome sequencing and there is a need to adapt short read tools to long read datasets.

**Methods:** We describe a new LCA-based algorithm for taxonomic binning, and an interval-tree based algorithm for functional binning, that are explicitly designed for long reads and assembled contigs. We provide a new interactive tool for investigating the alignment of long reads against reference sequences. For taxonomic and functional binning, we propose to use LAST to compare long reads against the NCBI-nr protein reference database so as to obtain frame-shift aware alignments, and then to process the results using our new methods.

**Results:** All presented methods are implemented in the open source edition of MEGAN and we refer to this new extension as MEGAN-LR (MEGAN long read). We evaluate the LAST+MEGAN-LR approach in a simulation study, and on a number of mock community datasets consisting of Nanopore reads, PacBio reads and assembled PacBio reads. We also illustrate the practical application on a Nanopore dataset that we sequenced from an anammox bio-rector community.

## Background

There are numerous computational tools for taxonomic or functional binning or profiling of microbiome samples, optimized to run on hundreds of millions of short, high quality sequencing reads [1, 2, 3, 4]. Alignment-based taxonomic binning is often performed using the naïve LCA algorithm [5], because it is fast, easy to interpret and easy to implement. Functional binning usually involves a best-hit strategy to assign reads to functional classes.

Software or websites for analyzing microbiome shotgun sequencing samples usually provide some level of interactivity, such as MG-RAST [2]. The interactive microbiome analysis tool MEGAN, which was first used in 2006 [6], is one of the most feature-rich tools of this type. MEGAN is highly optimized to enable users to interactively explore large numbers of microbiome samples containing hundreds of millions of short reads.

Illumina HiSeq and MiSeq sequencers allow researchers to generate sequencing data on a huge scale, so as to analyze many samples at a great sequencing depth [7, 8, 9]. A wide range of questions, in particular involving the presence or absence of particular organisms or genes in a sample, can be answered using such data. However, there are interesting problems that are not easily resolved using short reads. For example, the question whether two genes, which both are detected in the same microbiome *sample,* also occur together on the same *genome,* can often not be answered even for genes that lie close to each other in a genome, despite the use of contig binning techniques [10].

Current long read sequencing technologies produce smaller numbers (in the range of hundreds of thousands) of longer reads (of varying lengths in the range of 10 kb – 1 Mb, say) of lower quality (error rates around 10%, say). There is increasing interest in using long reads in microbiome sequencing and there is a need to adapt short read tools to long read datasets. There are a number of tools that are applicable to long reads, such as WIMP [11], Centrifuge [12] or Kaiju [13]. While the two former are based on comparing against DNA references, the latter can also use a protein reference database.

Here we present a new classification pipeline for taxonomic and functional analysis of long reads and contigs, based on protein alignments. The pipeline, LAST+MEGAN-LR, consists of first running the protein alignment tool LAST and then processing the resulting alignments using new algorithms provided in MEGAN-LR. We perform a simulation study to evaluate the performance of the method in the context of taxonomic assignment, and compare it with Kaiju, one of the few tools that works on protein references. We also investigate the performance of the pipeline using mock-community datasets and illustrate its use on Nanopore reads sequenced from an anammox enrichment bio-rector.

## Methods

### Long read taxonomic binning

The naïve LCA (lowest common ancestor) algorithm is widely used for binning short reads onto the nodes of a given taxonomy (such as the NCBI taxonomy), based on alignments [5]. Consider a read *r* that has significant alignments *a*_1_…, *a_k_* to reference sequences associated with taxa *t*_1_,…, *t_k_*. The naïve LCA assigns r to the lowest taxonomic node that lies above the set of all nodes representing *t*_1_,…, *t_k_*. The set of *significant* alignments is defined to consist of those alignments whose score lies close to the best score achieved for the given read, defined, say, as those that have a bit score that lies within 10% of the best bit score.

The naïve LCA algorithm is extremely fast, easy to implement and the results are easy to interpret. When applied to protein alignments, an implicit assumption of the algorithm is that any read aligns to only one gene and so all associated taxa are “competing” for the same gene; this justifies the above definition of significant alignments. While reads that are only a few hundred base pairs long usually fulfill this assumption, longer reads or assembled contigs often overlap with more than one gene and so the naïve algorithm is not suitable for them.

To make the naïve algorithm applicable to protein alignments on a long read or contig *r*, a simple idea is to first detect different “conserved genes” along the read as regions where alignments accumulate, and then to apply the naive LCA to each of these regions individually. The final placement of the read is then determined, for example, using the LCA of all the gene-based LCAs. There are two problems here. First, protein alignments around the same location can have quite different lengths and thus delineating different “conserved genes” can be difficult in practice. Second, because a large proportion of genes in a long read or contig may be conserved across different taxonomic groups, the final placement of the read will often be very unspecific.

To address these issues, we present a new taxonomic binning for long reads that we call the *interval-union LCA.* This algorithm processes each read *r* in turn, in two steps. First, the read is partitioned into a set of intervals *u*_1_,…, *υ_m_* that have the property that every alignment associated with r starts and ends at the beginning or end of some interval, respectively. In other words, a new interval starts wherever some alignment begins or ends. We say that an alignment *a_i_* is *significant* on an interval *υ_j_*, if its bit score lies within 10% (by default) of the best bit score seen for any alignment that covers *υ_j_*.

In the second step, for each taxon t that is associated with any of the alignments, let *I(t)* denote the union of all intervals for which there exists some significant alignment ¾ associated with taxon *t*. In a post-order traversal, for each higher-rank taxonomic node s we compute *I(s)* as the union of the intervals covered by the children of *s*. In result, every node of the taxonomy is labeled by a set of intervals. (Note that, during the computation of the union of interval sets, we merge any overlapping intervals into a single interval.)

The read *r* is then placed on the taxon *s* that has the property that its set of intervals *I*(*s*) covers 80% (by default) of the total aligned or covered portion of the read, while none of its children does (see Figure 1). Note that it theoretically possible that multiple nodes have this property, in which case the read is assigned to the LCA of all such nodes.

**Figure 1:**
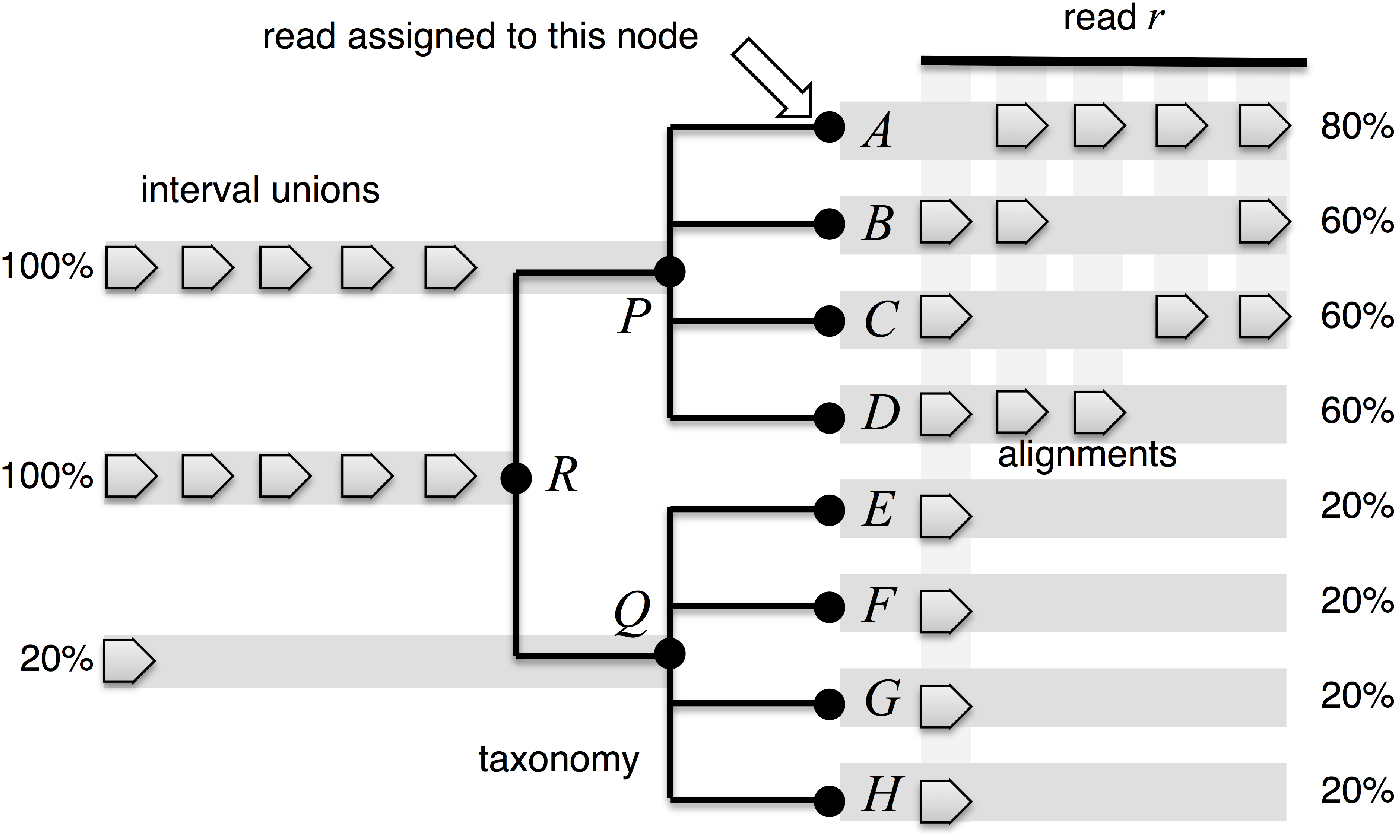
To illustrate the interval-union LCA algorithm, here we show eight hypothetical species *A,… H* separated into two genera, *P* and *Q*, belonging to a family *R*. Alignments from the read *r* to proteins associated with the species are indicated by arrows on the right and cover between 80% (for *A*) and 20% (for *H*) of the aligned read. Using arrows, on the left we depict the sets of intervals computed for nodes *P, Q, R* as the union of the sets of intervals of the children of each node. Nodes *R* and *P* each cover 100% of the aligned read. The read *r* is placed on *A* as it is the lowest taxonomic node with ≥ 80% coverage. Note that, if *A* only covered 60% of the aligned read, say, then the read would be assigned to taxon *P*, even if one of the taxa below *Q* also had 60% coverage, say.

### Long read functional binning and annotation

Functional binning of short reads is usually performed by assigning each read to a class in a functional classification system such as InterPro [14], eggNOG [15] or KEGG [16], based on its alignments.

This is often done using a simple *best-hit* strategy, as follows. For a short read *r*, let *a* denote the highest-scoring alignment of *r* to a reference protein for which the functional class *c* is known. Assign *r* to the functional class *c*. For example, *c* might be an InterPro family or an eggNOG cluster. In short read analysis, any read is assigned to at most one class in any given functional classification (and many reads remain unclassified because all corresponding reference proteins are unclassified.)

A long read may contain multiple genes and alignments will pile up at the locations of such genes. A functional binning algorithm for long reads and contigs must select a subset of alignments to report functional classes on, to avoid a high level of redundant or conflicting reports along the read.

Let *r* be a long read and let *a*_1_,…,*a_k_* be a set of DNA-to-protein alignments from *r* to a suitable protein reference sequences. For the sake of concreteness, let us assume that we are using the InterPro classification and that the InterPro family is known for each of the reference sequences.

To reduce the number of redundant functional classes associated with *r*, we introduce the following concept. We say that an alignment *a_i_ dominates* an alignment *a_j_*, if (1) *a_i_* covers more than 50% of the read that is covered by *a_j_*, (2) if the bit score of *a_i_* is greater than that of *a_j_*, and (3), both alignments lie on the same strand of *r*. Optionally, one could also require that the taxonomic identify of each protein reference sequence under consideration is compatible with the taxonomic bin assigned to the read *r*.

The set of functional classes associated with a long read r is then given by the functional classes associated with those alignments or *r* that are not dominated by some other alignment of *r*. Each read can be binned to all functional classes associated with it. Moreover, the set of associated classes can be used to provide a simple functional annotation of the read or contig.

To exploit that latter, we provide a dialog for exporting taxonomic and functional annotations in GFF3 format. It can be applied to any selection of taxonomic or functional classification nodes, or it can be applied to a set of selected reads in the new *long read inspector,* which is described in more detail below. The user chooses a classification and then each alignment to a reference sequence associated with that classification is exported as a CDS item. By default, only those alignments that are not dominated by some other alignment are exported. In addition, the user can decide to export only those items for which the taxon associated with the corresponding reference sequence is compatible with the taxon assigned to the read.

### Reporting counts

In taxonomic or functional binning of short reads, it usually suffices to report the number of reads assigned to a specific classification node, because all reads are of very similar length and all alignments have much the same length as the reads. For long reads or contigs, the lengths and alignment coverage can vary widely. Moreover, the number of reads contained in a contig, or contig coverage, is an additional factor to be considered. To address this, in MEGAN-LR each node can be labeled by one of the following:

1. the number of reads assigned,
2. the total length of all reads assigned,
3. the total number of aligned bases of all reads assigned, or
4. in the case of contigs, the total number of reads contained in all assigned contigs.

For long reads, by default, in MEGAN–LR report (3), the number of aligned bases, as this down-weights any long stretches of unaligned sequence. In addition, we use this value to determine the minimum support required for a taxon so that it is reported. By default, a taxon is only reported if it obtains at least 0. 05% of all aligned bases. If the number of aligned bases assigned to a taxon t does not meet this threshold, then the assigned bases are pushed up the taxonomy until a taxon is reached that has enough aligned bases to be reported.

### Long read alignment

In this paper we focus on taxonomic and functional binning of long reads using protein alignments. Currently long read sequencing technologies (Oxford Nanopore and PacBio) exhibit high rates of erroneous insertions and deletions. Programs such as BLASTX [17] and DIAMOND [18] cannot handle frame–shifts and so are not suitable for such reads.

The LAST program [19, 20] uses a frame–shift aware algorithm to align DNA to proteins and produces long protein alignments on long reads, even in the presence of many frame-shifts. Initial indexing of the NCBI-nr database (containing over 100 million sequences) by LAST takes over one day on a server. However, once completed, alignment of reads against the NCBI-nr database using the index is fast; the alignment of Nanopore reads takes roughly one hour per gigabase on a server.

### Long read analysis

LAST produces output in a simple text–based multiple alignment format (MAF). For performance reasons, LAST processes all queries and all reference sequences in batches and in consequence, alignments associated with a given query are not reported consecutively, but rather in batches.

In addition, the size of a MAF file is often very large and subsequent sorting and parsing of alignments can be time consuming. To address these issues, we have implemented a new program called “MAF2DAA” that takes MAF format as input, either as a file or piped directly from LAST, and produces a DAA file as output. The program processes the input in chunks, first filtering and compressing each chunk of data on-the-fly, and then interleaving and filtering the results into a single DAA file that contains all reads with their associated alignments. During filtering, MAF2DAA removes all alignments that are *strongly dominated* by some other alignment, so as to reduce a large amount of redundant alignments.

In more detail, for a given read *r*, we say that an alignment *a* of *r strongly dominates* an alignment *b* for *r*, if it covers most of *b* (by default, we require 90% coverage) and if its bit score is significantly larger (by default, we require that 0.9 · bitscore(*a*) > bitscore(*b*)).

A DAA file obtained in this way can then be processed by MEGAN’s Meganizer program that performs taxonomic and functional binning, and indexing, of all reads in the DAA file. This program does not produce a new file, but rather simply appends the results to the end of the DAA file and any such “meganized” DAA file can be directly opened in MEGAN for interactive analysis. We have modified MEGAN so that it supports frame-shift containing alignments. The final DAA file is usually around 10 times smaller than the MAF file produced by LAST.

### Long read visualization

Interactive analysis tools for short read microbiome sequencing data usually focus on representing the taxonomic and functional classifications systems used for binning or profiling the reads, for example reporting the number of reads assigned to each class. In addition, some tools provide a reference-centric visualization that displays how the reads align against a given reference sequence. However, visualizations of the short reads themselves are usually not provided.

For long read or contigs, there is a need for visualization techniques that make it easy to explore the taxonomic and functional identity of reference sequences to which the reads align. To address this, we have designed and implemented a *long read inspector* (using JavaFX) that allows one to investigate all long reads assigned to a given taxonomic or functional class (see Figure 2).

**Figure 2:**
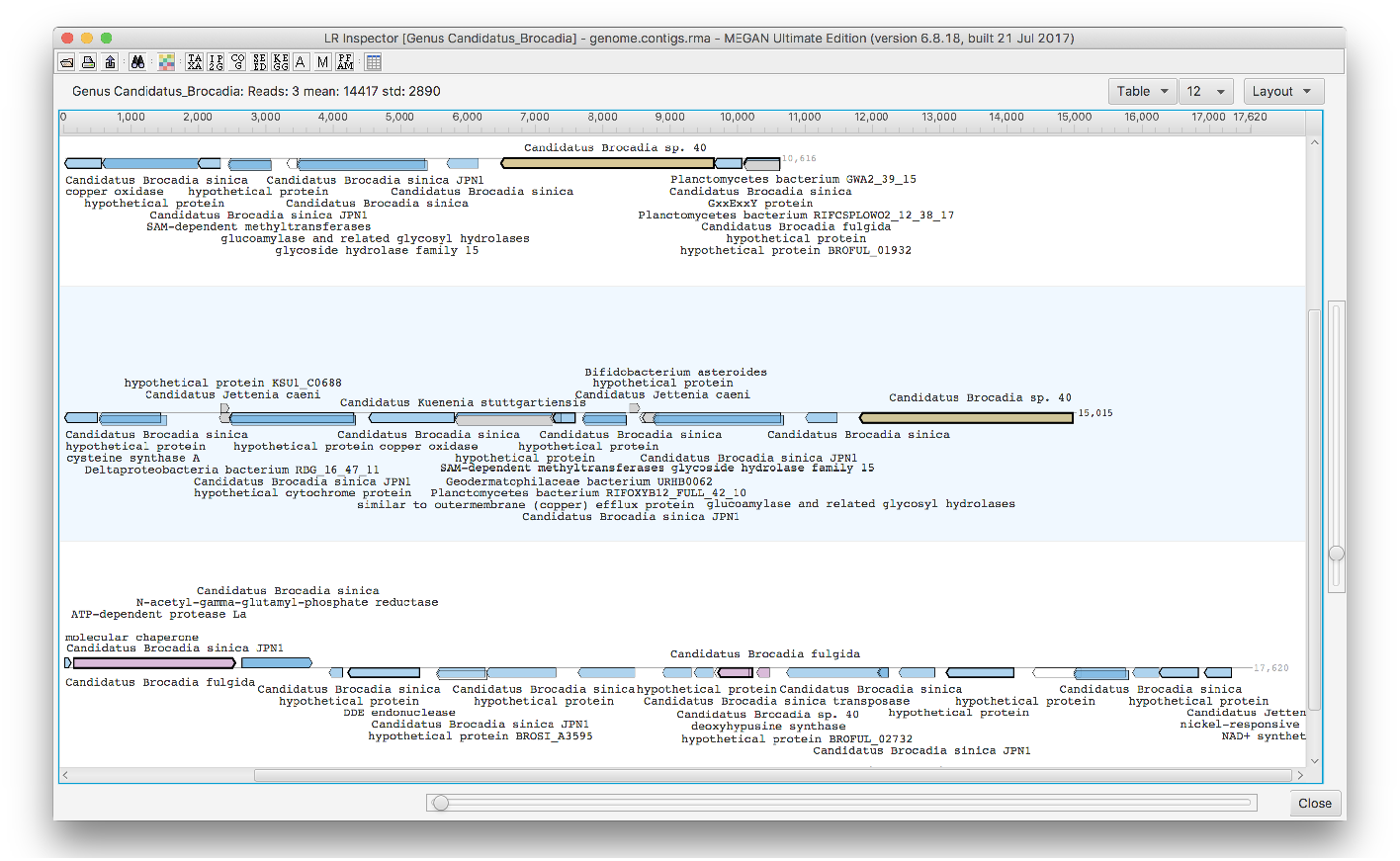
This screen shot of the MEGAN-LR long read inspector shows three contigs assigned to the genus *Candidatus Brocadia.* Alignments to reference protein sequences are shown as arrows, colored by species of the references; blue for *Candidatus Brocadia sinica,* brown for *Candidatus Brocadia sp. 40* and pink for *Candidatus Brocadia fulgida.* Alignments are labeled by taxonomic and functional classes associated with the corresponding reference proteins.

In this tool, each long read or contig r is represented by a horizontal line and all corresponding aligned reference sequences are shown as arrows above (forward strand alignments) or below (reverse strand alignments) the line. The user can select which annotations to display in the view. For example, if the the user requests Taxonomy and InterPro annotations, then all reference sequences will be labeled by the associated taxonomic and InterPro classes. The user can search for functional attributes in all loaded reads.

Let *a* be an arrow representing an alignment of r to a reference sequence associated with taxon *s*. We use a hierarchical coloring scheme to color such arrows. Initially, we implicitly assign a color index to each taxon, e.g. using the hash code of the taxon name. For each arrow *a* with associated reference taxon *s* we distinguish between three different cases. First, if *s* = *t*, then we use the color assigned to *t* to color *a*. Second, if *s* is a descendant of *t*, then *t* has a unique child *u* that lies on the path from *t* down to *s* and we use the color of *u* to color *a*. Otherwise, we color *a* gray to indicate that the taxon associated with *a* is either less specific or incompatible with *t*.

For example, if a read *r* is assigned to the genus *Candidatus Brocadia* and has an alignment to the strain *Candidatus Brocadia sinica JPN1,* then we color the corresponding arrow *a* using the color that represents the species *Candidatus Brocadia sinica.*

This is a useful strategy when used in combination with the taxonomic binning procedure described above: *a* read *r* is binned to the lowest taxon *t* that covers 80% (by default) of the aligned read and the taxonomy-based coloring makes it easy to see how the different taxonomic classes below *t* contribute. For example, if all arrows on one half of the read have one color and all arrows on the other half have some other color, then this may indicate a chimeric read or misassembled contig.

As discussed above, an alternative approach is to export reads and their alignments in GFF format and then to use a genome browser such as IGB [21] to explore them (see Figure 3).

**Figure 3:**
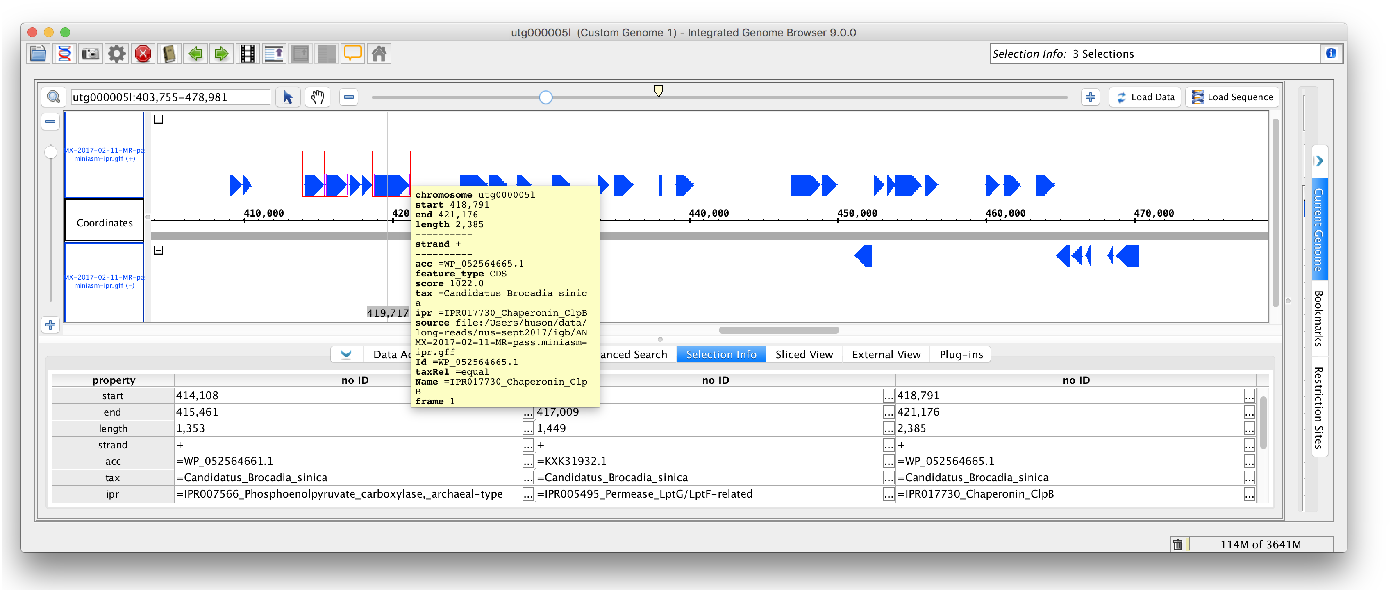
Example of long read data exported from MEGAN-LR and imported into the IGB genome browser.

## LAST+MEGAN-LR

In summary, we propose to use the following pipeline to analyze metagenomic long reads and contigs (see Figure 4):

- Align all reads against a protein reference database (such as NCBI-nr) using LAST, producing MAF output.
- Either pipe the output of LAST directly to MAF2DAA, or apply MAF2DAA to the MAF file generated by LAST, so as to obtain a much smaller output file in DAA format.
- Meganize the DAA file either using the Meganizer command-line tool, or interactively in MEGAN.
- Open the meganized DAA file in MEGAN for interactive exploration using the long-read inspector. Export annotated reads in GFF format for further investigation, e.g. using a genome browser such as Artemis [22] or IGB [21].

**Figure 4:**
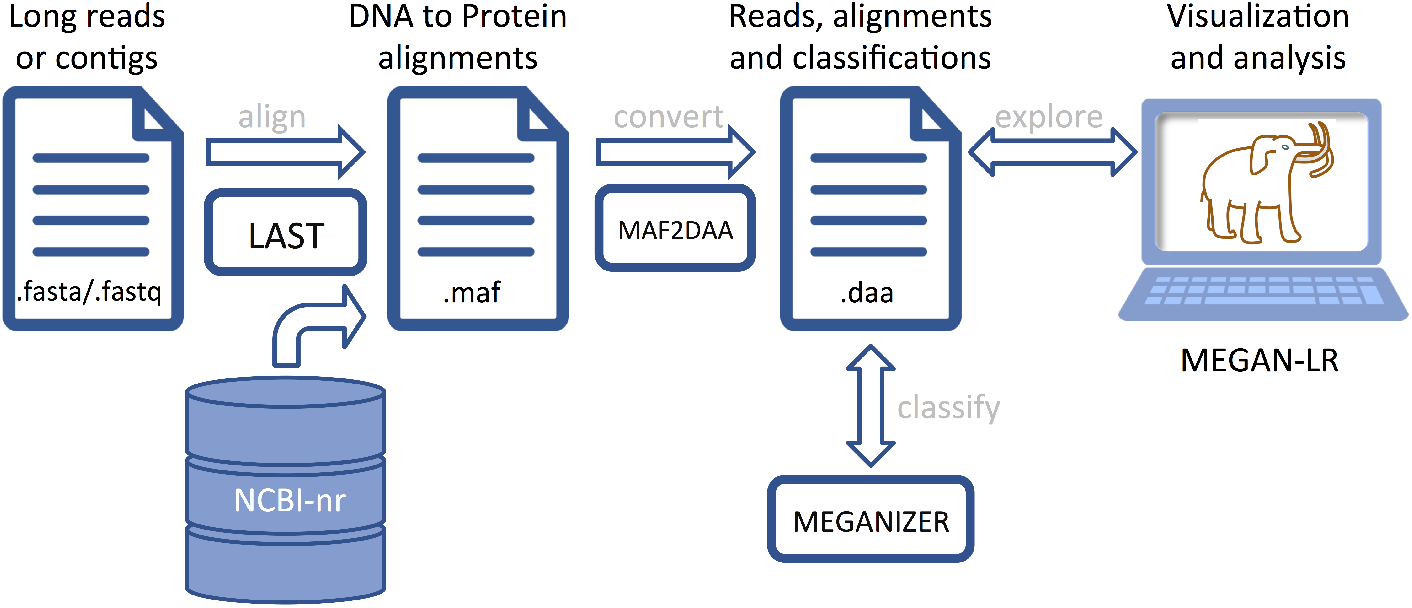
The LAST+MEGAN-LR pipeline. Long reads or contigs are aligned against the NCBI-nr database using LAST and the resulting MAF file (multiple alignment format) is converted to DAA format (Diamond alignment format), including filtering of dominated alignments. Taxonomic and functional binning of the reads or contigs is then performed using the Meganizer program and the results are stored to the DAA file. The meganized DAA file can then opened and analyzed in MEGAN-LR.

### Nanopore Sequencing

To obtain a Nanopore sequencing dataset for method evaluation, we sequenced the genomic DNA of the Microbial Mock Community B (even, high concentration, catalog nr. HM-276D, BEI Resources). Library preparation was performed using a Low Input by PCR Genomic Sequencing Kit SQK-MAP006 (Oxford Nanopore Technologies, Oxford, UK) for 2D sequencing. Briefly, 100 ng of genomic DNA was sheared in a Covaris g-TUBE (Covaris, Inc., Woburn, MA, USA) at 6000 rpm, treated with PreCR (New England Biolabs, Ipswich, MA, USA) and used as input for adapter ligation according to the ONT protocol. Adapter-ligated DNA was further amplified with the LongAmp Taq 2X Master Mix (NEB) using the following program: 95°C 3 min; 18 cycles of 95°C 15 sec, 62°C 15 sec, 65°C 10 min; 65°C 20 min. Sequencing was performed using an early access MinION device (ONT) on a FLO-MAP003 flowcell (ONT). Raw fast5 files were obtained with MinKNOW (v0.50.2.15, ONT) using a 48 h genomic sequencing protocol, basecalled with ONT’s proprietary Metrichor cloud-based basecalling service and the 2D Basecalling for SQK-MAP006 v1.34 workflow.

Genomic DNA from the lab scale Anammox enrichment reactor described in Liu *et al.* [23] was extracted using FastDNA SPIN Kit for Soil with 4x homogenization on the FastPrep instrument (MP Bio). The DNA was further purified using Genomic DNA Clean and Concentator −10 Kit (Zymo Research). ≈ 1700 ng of extracted DNA was used for library preparation using a Ligation Sequencing Kit SQK-LSK108 (Oxford Nanopore Technologies, Oxford, UK) for 1D sequencing according to the manufacturer protocol. Sequencing was performed using an early access MinION device (ONT) on a SpotON FLO-MIN106 flowcell (R9.4). The run was stopped after 22 hours due to low number of active pores. Fast5 files were obtained with MinKNOW (v1.3.30, ONT) using a 48 h genomic sequencing protocol. Basecalling was performed using Metrichor (Instance ID:135935, 1D Basecalling for FLO-MIN106 450 bps_RNN (rev.1.121)).

### Simulation study

To evaluate the performance of the proposed LAST+MEGAN-LR approach and, in particular, of the interval-union LCA algorithm, we undertook a simulation study to estimate the sensitivity and precision of the algorithm, following the protocol reported in [13] We attempted to model two major obstacles in metagenomic studies, namely sequencing errors and the incompleteness of reference databases.

Our simulation study is based on a set *P* of 4, 282 prokaryotic genomes from NCBI for which both annotated genomes and annotated sets of proteins are available, downloaded in March 2017. In addition, we defined a subset *Q* of 1,151 genomes that consists of all those organisms in *P* whose genus contains at least 2 and at most 10 organisms in *P*, and for which a full taxonomic classification is given. Notice that *Q* can be partitioned into nine different categories, based on the number 2 – 10 of organisms in *Q* that the corresponding genus contains.

For each target species *t* in *Q*, we performed the following leave-one-out evaluation:

- First, we collected a set of *R* of 1,000 simulated reads from the genome sequence of *t* using NanoSim [24], a read simulator that produces synthetic reads that reflect the characteristic base-calling errors of ONT reads, running in chromosomal circular mode.
- Second, we constructed a protein reference database 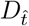 that contained all annotated proteins associated with all organisms in *P* except for *t*.
- Third, we performed taxonomic binning of all reads in *R* using LAST+MEGAN-LR as follows. We first build a LAST reference index on 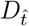, then aligned all reads in *R* against 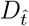 using LAST with a frameshift cost of 15, and then performed taxonomic binning of all reads in MEGAN using the interval-union LCA algorithm (default parameters).
- Fourth, for comparison, we also ran the taxonomic binning program Kaiju[13] on *R* and 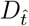, as this is one the few tools that performs taxonomic binning using a protein reference database. We built custom Kaiju indexes of 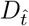. We performed taxonomic binning of simulated reads using Kaiju’s greedy mode, with the maximum number of allowed substitutions set to 5.

To be precise, we ran each of the four steps twice so as to produce two simulation datasets, each containing 1,000 reads per target species. The first dataset was produced using the ecoli_R73_2D (R7.3) simulator profile, whereas the second was produced using the ecoli_R9_2D (R9) profile. Both profiles were downloaded from the NanoSim FTP address (ftp://ftp.bcgsc.ca/supplementary/NanoSim/) in April 2017. The R7.3 profile introduces more errors in reads and should make it harder for analysis methods to identify appropriate reference sequences.

To compare the performance of MEGAN-LR and Kaiju, we calculated the sensitivity and precision with which both programs made taxonomic assignments at the genus, family and order levels. In more detail, following the approach used in [13], we define *sensitivity* as the percentage of reads in R that are assigned to the correct taxon. We define *precision* as the percentage of reads that are assigned correctly, out of all reads that were binned to any node that is not an ancestor of the correct taxon.

## Results

We have implemented the interval-union taxonomic binning algorithm and modified functional binning algorithms. In addition, we have implemented a new long read interactive viewer (using JavaFX). We provide methods for exporting long read annotations in GFF3 format. Our code has been integrated into the open source edition of MEGAN. In addition, we have modified MEGAN (and all tools bundled with MEGAN) so as to support DNA-to-protein alignments that contain frame-shifts. We use the term MEGAN-LR (MEGAN long read) to refer to this major extension of MEGAN.

### Simulation study

The results of our simulation study are shown in Figure 5, where we summarize the sensitivity and precision scores achieved at genus level by LAST+MEGAN-LR and Kaiju, for both the R7.3 and R9 datasets. In all cases, LAST+MEGAN-LR shows better sensitivity and precision than Kaiju. As expected, both methods are less sensitive on the R7.3 data, as many reads remain unclassified. However, the difference in performance between the two methods is larger on the R7.3 data and we suspect that this is due to the ability of LAST to perform frame-shift aware alignments and thus to accommodate erroneous insertions and deletions.

**Figure 5:**
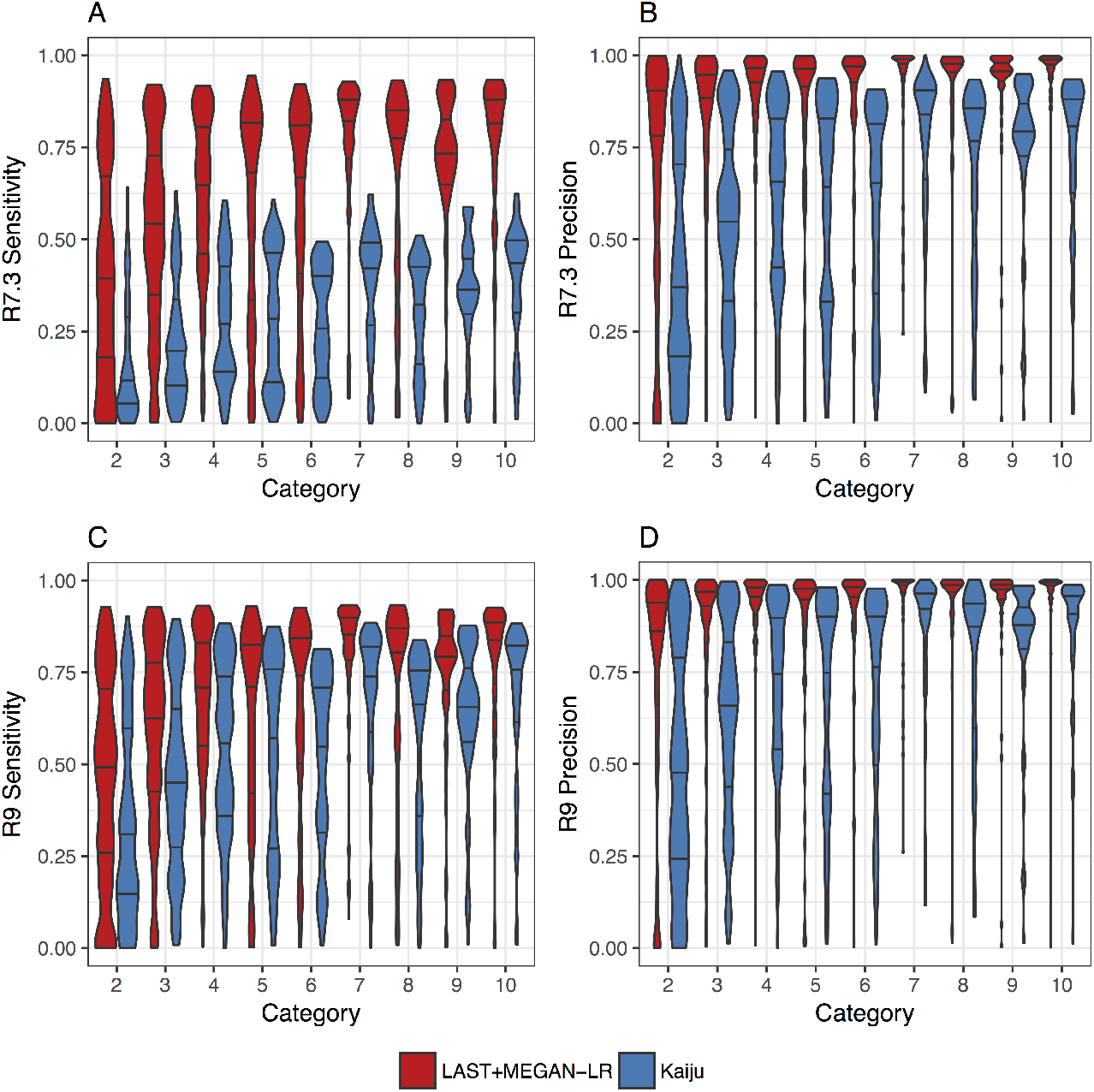
Simulation study results for two simulation studies based on the R7.3 and R9 Nanopore chemistry profiles. We report the sensitivity and precision of LAST+MEGAN-LR and Kaiju at genus level for 9 categories (reflecting the number of species in the genus from which the target species was removed). We report the sensitivity and precision for the R7.3 profile in (A) and (B), respectively, and for the R9 profile in (C) and (D), respectively.

A per-dataset performance analysis of LAST+MEGAN-LR and Kaiju is presented in Figure 6. This shows that LAST+MEGAN-LR outperforms Kajiu on a vast majority of the simulated datasets, with Kajiu sometimes showing better performance at when the sensitivity or precision is very low.

**Figure 6:**
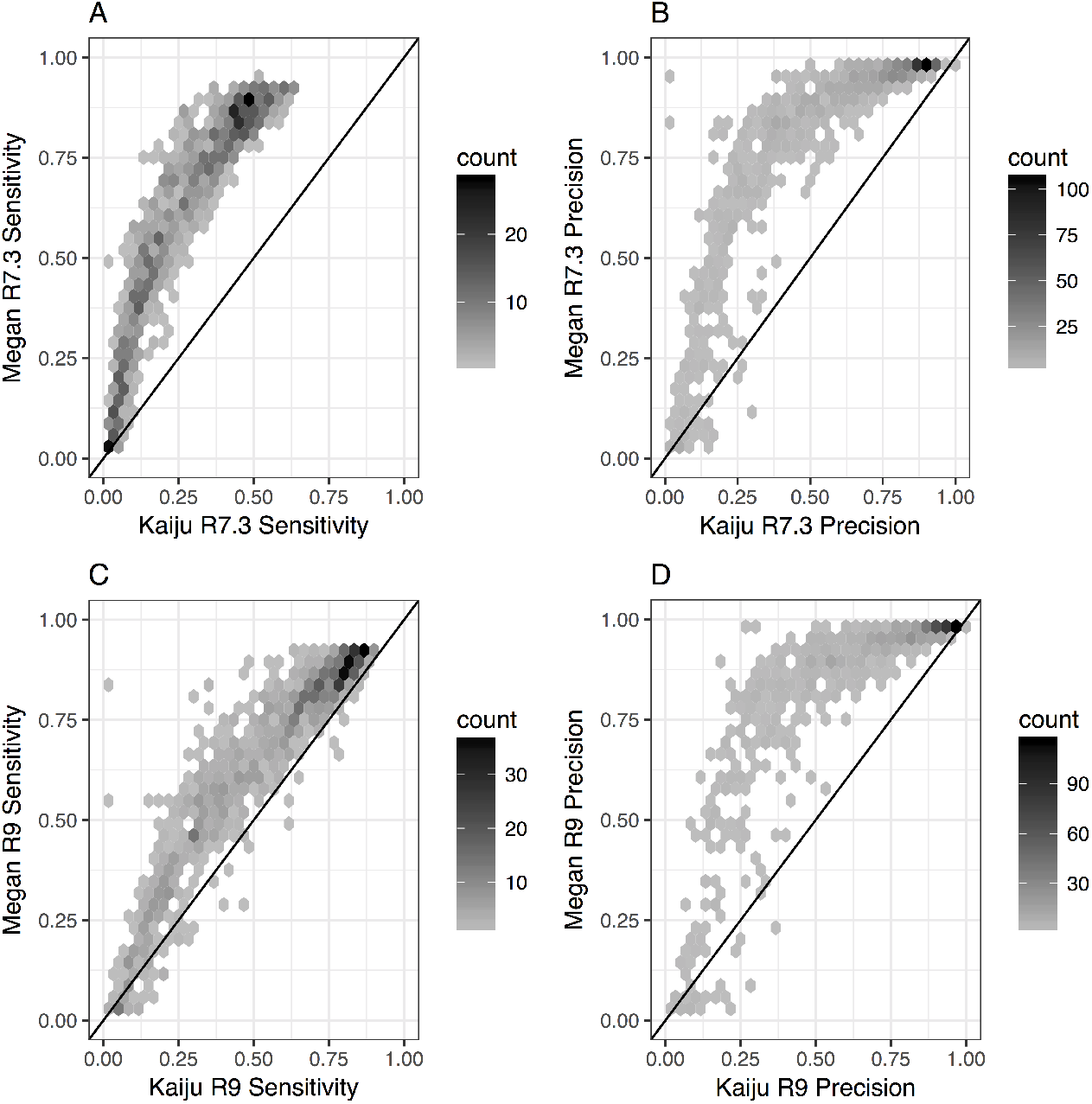
Here we plot the sensitivity and precision at genus level for Kaiju versus LAST+MEGAN-LR on the R7.3 samples in (A) and (B), respectively, and on the R9 samples in (C) and (D), respectively.

Kaiju is many times faster than LAST+MEGAN-LR. However, the latter approach computes and uses all relevant protein alignments, and these are also used to perform functional analysis of the reads or contigs. Hence, we suggest to use Kaiju to obtain a fast, first taxonomic profile for a set of long reads or contigs, and then to use LAST+MEGAN-LR to perform a more accurate and detailed subsequent analysis.

### PacBio reads on HMP mock community

To test LAST+MEGAN-LR on a publicly available PacBio mock community dataset, we downloaded “HMP dataset 7” from the PacBio website https://github.com/PacificBiosciences/DevNet/wiki/Human_Microbiome_Project_MockB_Shotgun in April 2017. This dataset contains 319, 703 reads of average length 4, 681 bp. It was sequenced using the P5 polymerase and C3 chemistry.

LAST alignment against the NCBI-nr database (downloaded January 2017) resulting in protein alignments for 279, 926 reads (87% of all reads). MEGAN-LR analysis using the interval-union assigned 1, 000 megabases (Mb) aligned bases to taxonomic nodes. Of these, 923.8 Mb were assigned to bacterial genera, with a false positive rate of 0.2%. A total of 743.6 Mb of aligned sequences were assigned to bacterial species, of which 741.6 Mb were assigned to true positive species (that is, species known to be contained in the mock-community), whereas approximately 2 Mb were assigned to false positive species, resulting in a false positive binning rate of 0.3%. The 20 bacterial species in the mock community received between 2.6 Mb (0.3%) and 143 Mb (19%) aligned bases assigned at the species level, whereas the highest false positive species obtained 1.3 Mb (0.2%).

To investigate the use of LAST+MEGAN-LR on assembled reads, we assembled this set of reads using minimap (options −Sw5 −L100 −m0 −t8) and miniasm (default options) [25] and obtained 1,130 contigs, with a mean length of 43, 976 and maximum length of 1, 272, 994. LAST alignment against the NCBI-nr database resulted in 41.8 Mb of aligned sequences. Of this, 40 Mb and 37.3 Mb, were assigned to genus and species nodes, respectively, with no false positives and only one false negative species.

### PacBio reads on another mock community

Analysis of PacBio reads recently published on a mock-community containing 26 bacterial and archaeal species [26] gave rise to similar results. Of 53, 654 reads of average length 1, 041 and maximum length 16,403, exactly 51,124 received LAST alignments against NCBI-nr. Of 49.4 Mb of aligned sequences, 45.6 Mb were assigned to prokaryotic genera, with no assignments to false positive species.

The amount of sequence assigned at the species level was 36.8 Mb, all of which was assigned to true positive species.

Of the 26 species in the mock community, there are two that are not reported in the analysis and therefore constitute false negative species. These make up approximately 0.01% (*Nocardiopsis dassonvillei*) and 0.1% (*Salmonella bongori*) of the community and are thus on the borderline of detection using the default settings of MEGAN-LR. By default, MEGAN-LR requires that a taxon receives at least 0.05% of all aligned bases before it is reported.

### Nanopore reads on HMP mock community

To perform a first test of our new methods on Nanopore data, we sequenced the content of the Genomic DNA from Microbial Mock Community B, as described in the Methods section. We obtained 124, 911 pass reads of average length 2, 870, including all template, complement and 2D reads.

LAST alignment against the NCBI-nr database resulting in protein alignments for 61, 261 reads (49% of all reads). MEGAN-LR analysis assigned a total of 110.8 Mb aligned bases. Of these, 96.8 MB were assigned to bacterial genera, with a false positive assignment of 0.3%. Approximately 71.2 Mb of aligned sequences were assigned at the species level, with a false positive rate of 0.6%. The 20 bacterial species in the mock community received between 0.3 Mb (0.5%) and 12 Mb (17%) aligned bases assigned at the species level, whereas the highest false positive species obtained 0.2 Mb (0.3%). Around 60 kb of all aligned sequences (0.05%) were falsely assigned to Eukaryota.

### Application to anammox data

To illustrate the utility of our new methods in a research context, we applied Nanopore sequencing to a sample obtained from a laboratory bio-reactor enriched for anaerobic ammonium oxidizing bacteria (AnAOB) [27], as described in the Methods section. We obtained 71, 411 reads of average length 4, 658 and maximum length 30, 846.

LAST alignment against the NCBI-nr database resulted in protein alignments for 64,097 reads (90% of all reads). MEGAN-LR analysis assigned a total of 209 Mb aligned bases. Of these, 96.3 Mb were assigned to bacterial genera and 113 Mb to bacterial species. The reason why there are more assignments to species than there are to genera is that a number of the species are not contained in any genus. The top ten bacterial species assignments are shown in Table 1. This indicates that the most abundant organism in the sample is *Candidatus Brocadia sinica,* a known AnAOB species.

**Table 1:** The ten top bacterial species identified in a Nanopore dataset taken from an anammox enrichment bioreactor. The number of aligned bases assigned to *Candidatus Brocadia sinica* suggest at least ten-fold coverage of that genome.

| Species | Assigned bases |
| --- | --- |
| *Candidatus Brocadia sinica* | 85.9 Mb |
| *Armatimonadetes bacterium OLB18* | 9.1 Mb |
| *Bacteroidetes bacterium OLB12* | 5.0 Mb |
| *Chloroflexi bacterium OLB13* | 4.8 Mb |
| *Nitrospira sp. OLB3* | 1.5 Mb |
| *Streptomyces sp. SolWspMP-5a-2* | 1.5 Mb |
| *Candidatus Brocadia sp. 40* | 0.66 Mb |
| *Burkholderia pseudomallei* | 0.54 Mb |
| *Chlorobi bacterium OLB4* | 0.49 Mb |
| *Chlorobi bacterium OLB15* | 0.48 Mb |

To illustrate the use of LAST+MEGAN-LR on assembled reads, we assembled this set of reads using minimap (options −Sw5 −L100 −m0 −t8) and miniasm (default options) [25] and obtained 31 contigs, with a mean length of 129, 601 and maximum length of 750, 799. LAST alignment against the NCBI-nr database resulted in 2.98 Mb of aligned sequences. The interval-union algorithm assigned 13 reads and 96% of all aligned bases to *Candidatus Brocadia sinica*. Another 6 reads and 1.5% of all aligned were assigned to *Armatimonadetes bacterium OLB18.*

## Discussion

The application of long read sequencing technologies to microbiome samples promises to provide a much more accurate description of the genomes present in environmental samples. The alignment of long reads against a protein reference database is a key step in the functional analysis of such data. Here we show that such protein alignments can also be used to perform accurate taxonomic binning using the interval-union LCA algorithm.

Our simulation study suggests that using LAST+MEGAN-LR performs taxonomic binning of long reads accurately, even in the absence of the same species in the reference database. The reported results on mock community datasets suggest a high level of accuracy down to the species level, when the corresponding species are represented in the protein reference database. In addition, the computed protein alignments can be used to identify genes and MEGAN-LR provides a useful visualization of the annotated sequences.

A main motivation for developing these new methods was to assist our work on the study of micro bial communities in enrichment bio-rectors, where long read sequencing promises to provide access to near complete genome sequences of the dominating species.

The simple assembly of the anammox data presented in this paper places the dominant species into 11 contigs of length greater than 100 kb, containing about 2.8 Mb of aligned sequence and 3.7 Mb of total sequence. This suggests that a more careful assembly, assisted by a set of high quality MiSeq reads, should result in a near complete genome.

Our simulation study indicates that LAST+MEGAN-LR is much more accurate than Kaiju. Because Kaiju uses a heuristic based on the longest match found, we suspect that Kaiju will perform poorly on chimeric reads or misassembled contigs, assigning such a read to one of the source taxa. In contrast, the interval-union requires by default that 80% of the aligned read is assigned to a taxon and so in practice, such reads will often be placed on a higher taxonomic node.

All datasets discussed in this paper are available here: http://ab.inf.uni-tuebingen.de/software/downloads/megan-lr

**List of abbreviations**
MEGAN-LR: long read extension of the metagenome analysis tool MEGAN

## Ethics approval and consent to participate

Not applicable.

## Consent for publication

Not applicable.

## Availability of data and materials Competing interests

The authors declare that they have no competing interests.

## Author’s contributions

All authors contributed to the study design. AG, DH and RW designed the visualization techniques. DH implemented the algorithms and visualization techniques. IB and DJ performed Nanopore sequencing. BA implemented the MAF2DAA program. BA and CB performed the simulation study. DH and RW lead the project and wrote the manuscript. All authors read and approved the final manuscript.

## Acknowledgements

The authors acknowledge support by the High Performance and Cloud Computing Group at the Zentrum für Datenverarbeitung of the University of Tübingen, the state of Baden-Württemberg through bwHPC and the German Research Foundation (DFG) through grant no INST 37/935-1 FUGG. This was supported in part by the Singapore National Research Foundation and Ministry of Education under the Research Centre of Excellence Programme, and by a program grant from the Environment and Water Industry Programme Office (EWI), project number 1301–IRIS–59. We thank Dr Gayathri Natarajan and Dr Ying Yu Law for their assistance with obtaining samples from the Anammox bioreactor.

## References

[1] D. Huson, S. Beier, I. Flade, A. Górska, M. El-Hadidi, S. Mitra, H.-J. Ruscheweyh, and R. Tappu. MEGAN Community Edition - interactive exploration and analysis of large-scale microbiome sequencing data. PLoS Comput Biol, 12(6):e1004957, 2016. doi:10.1371/journal.pcbi.1004957.

[2] E. M. Glass, J. Wilkening, A. Wilke, D. Antonopoulos, and F. Meyer. Using the metagenomics RAST server (MG–RAST) for analyzing shotgun metagenomes. Cold Spring Harb Protoc., 2010(1):pdb.prot5368, 2010.

[3] D. E. Wood and S. L. Salzberg. Kraken: ultrafast metagenomic sequence classification using exact alignments. Genome Biology, 15:R46, 2014.

[4] N. Segata, L. Waldron, A. Ballarini, V. Narasimhan, O. Jousson, and C. Huttenhower. Metagenomic microbial community profiling using unique clade-specific marker genes. Nat Meth, 9(8):811–814, 2012.

[5] D. H. Huson, A. F. Auch, J. Qi, and S. C. Schuster. MEGAN analysis of metagenomic data. Genome Res, 17(3):377–386, March 2007.

[6] H. N. Poinar, C. Schwarz, J. Qi, B. Shapiro, R. D. E. Macphee, B. Buigues, A. Tikhonov, D. H. Huson, L. P. Tomsho, A. Auch, M. Rampp, W. Miller, and S. C. Schuster. Metagenomics to paleogenomics: large-scale sequencing of mammoth DNA. Science, 311(5759):392–394, 2006.

[7] R. Mackelprang, M. Waldrop, K. DeAngelis, M. David, K. Chavarria, S. Blazewicz, E. Rubin, and J. Jansson. Metagenomic analysis of a permafrost microbial community reveals a rapid response to thaw. Nature, 480(7377):368–371, 2011.

[8] The Human Microbiome Project Consortium. Structure, function and diversity of the healthy human microbiome. Nature, 486(7402):207–214, 2012.

[9] M. Willmann, M. El-Hadidi, D. Huson, M. S chütz, C. Weidenmaier, I. Autenrieth, and S. Peter. Antibiotic selection pressure determination through sequence-based metagenomics. Antimicrobial Agents and Chemotherapy, 59(12):7335–45, 2015.

[10] D. D. Kang, J. Froula, R. Egan, and Z. Wang. MetaBAT, an efficient tool for accurately reconstructing single genomes from complex microbial communities. PeerJ, 3:e1165, 2015.

[11] S. Juul, F. Izquierdo, A. Hurst, X. Dai, A. Wright, E. Kulesha, R. Pettett, and D. J. Turner. What’s in my pot? Real–time species identification on the MinION. bioRxiv, 030742, 2015.

[12] D. Kim, L. Song, F. P. Breitwieser, and S. L. Salzberg. Centrifuge: rapid and sensitive classification of metagenomic sequences. Genome Research, 26:1721–1729, 2016.

[13] P. Menzel, K. L. Ng, and A. Krogh. Fast and sensitive taxonomic classification for metagenomics with Kaiju. Nature Communications, 7:11257, 2016.

[14] A. Mitchell, H.-Y. Chang, L. Daugherty, M. Fraser, S. Hunter, R. Lopez, C. McAnulla, C. McMenamin, G. Nuka, S. Pesseat, A. Sangrador-Vegas, M. Scheremetjew, C. Rato, S.-Y. Yong, A. Bateman, M. Punta, T. K. Attwood, C. J. Sigrist, N. Redaschi, C. Rivoire, I. Xenarios, D. Kahn, D. Guyot, P. Bork, I. Letunic, J. Gough, M. Oates, D. Haft, H. Huang, D. A. Natale, C. H. Wu, C. Orengo, I. Sillitoe, H. Mi, P. D. Thomas, and R. D. Finn. The InterPro protein families database: the classification resource after 15 years. Nucleic Acids Research, 43 (Database Issue):D213–D221, 2015.

[15] S. Powell, D. Szklarczyk, K. Trachana, A. Roth, M. Kuhn, J. Muller, R. Arnold, T. Rattei, I. Letunic, T. Doerks, L. J. Jensen, C. von Mering, and P. Bork. eggNOG v3.0: orthologous groups covering 1133 organisms at 41 different taxonomic ranges. Nucleic Acids Research, 40(Database Issue):284–289, 2012.

[16] M. Kanehisa and S. Goto. KEGG: Kyoto Encyclopedia of Genes and Genomes. Nucleic Acids Res, 28(1):27–30, Jan 2000.

[17] S. Altschul, T. Madden, A. Schaffer, J. Zhang, Z. Zhang, W. Miller, and D. Lipman. Gapped BLAST and PSI-BLAST: a new generation of protein database search programs. Nucleic Acids Res., 25:3389–3402, 1997.

[18] B. Buchfink, C. Xie, and D. Huson. Fast and sensitive protein alignment using DIAMOND. Nature Methods, 12:59–60, 2015.

[19] S. M. Kielbasa, R. Wan, K. Sato, P. Horton, and M. C. Frith. Adaptive seeds tame genomic sequence comparison. Genome Research, 21(3):487–493, 2011.

[20] S. L. Sheetlin, Y. Park, M. C. Frith, and J. L. Spouge. Frameshift alignment: statistics and post-genomic applications. Bioinformatics, 30(24):3575–3582, 2014.

[21] J. W. Nicol, G. A. Helt, S. G. Blanchard, Jr., A. Raja, and A. E. Loraine. The Integrated Genome Browser: free software for distribution and exploration of genome-scale datasets. Bioinformatics, 25(20):2730–2731, 2009.

[22] K. Rutherford, J. Parkhill, J. Crook, T. Horsnell, P. Rice, M. A. Rajandream, and B. Barrell. Artemis: sequence visualization and annotation. Bioinformatics, 16(10):944–945, 2000.

[23] X. Liu, K. Arumugam, G. Natarajan, S. T. W., D. I. Drautz-Moses, S. Wuertz, L. Yingyu, and R. B. H. Williams. Draft genome sequence of a *Candidatus* brocadia bacterium enriched from tropical–climate activated sludge. BioRvix, 123943, 2017.

[24] C. Yang, J. Chu, R. L. Warren, and I. Birol. NanoSim: nanopore sequence read simulator based on statistical characterization. GigaScience, 6(4):1–6, 2017.

[25] H. Li. Minimap and miniasm: fast mapping and de novo assembly for noisy long sequences. Bioinformatics, 32(14):2103–2110, 2016.

[26] E. Singer, B. Andreopoulos, R. M. Bowers, J. Lee, S. Deshpande, J. Chiniquy, D. Ciobanu, H.-P. Klenk, M. Zane, C. Daum, A. Clum, J.-F. Cheng, A. Copeland, and T. Woyke. Next generation sequencing data of a defined microbial mock community. Scientific Data, 3:160081, 2016.

[27] B. Kartal, N. M. de Almeida, W. J. Maalcke, H. J. Op den Camp, M. S. Jetten, and J. T. Keltjens. How to make a living from anaerobic ammonium oxidation. FEMS Microbiology Reviews, 37:428–461, 2013.

